# An MILP Model for Corn Planting and Harvest Scheduling Considering Storage Capacity and Growing Degree Units

**DOI:** 10.1101/2021.02.06.430062

**Authors:** Zahra Khalilzadeh, Lizhi Wang

## Abstract

Corn planting and harvest scheduling is an important problem due to having a significant impact on corn yield, balancing the capacities for harvest, transport, and storage operations. Different corn hybrids also have different planting window and poor planting and harvest schedules may cause erratic weekly harvest quantities and logistical and productivity issues. In the 2021 Syngenta Crop Challenge, Syngenta released several large datasets that recorded the historical daily growing degree units (GDU) of two sites and provided planting window, required GDUs, and harvest quantity of corn hybrids planted in these two sites. Then, participants of this challenge were asked to schedule planting and harvesting dates of corn hybrids under two storage capacity scenarios so that facilities are not over capacity in harvesting weeks and have consistent weekly harvest quantities. The two storage capacity scenarios include: (1) planting and harvest scheduling given the maximum storage capacity, and (2) planting and harvest scheduling without maximum storage capacity to determine the lowest possible capacity for each site. In this paper, we propose two mixed integer linear programming (MILP) models for solving this problem considering both the storage capacity and the uncertainty in GDUs. Our results indicate that our proposed models can provide optimal planting and harvest scheduling under different GDU possibilities which ensures consistent weekly harvest quantities that are below the maximum capacity.

## 1 Introduction

The advent of new farming technologies such as commercial hybrid in the 1930s, preceded a widespread and rapid replacement of the once predominant open-pollinated seed varieties planted by farmers [1]. The widespread use of commercial hybrids is seen in many crops, including corn, sorghum, sugarbeet, and sunflower [2]. Of these crops, corn is widely known as one of the world’s most produced and important crops.

Scheduling planting and harvesting dates of the corn hybrids is an important part of corn crop production. Accurate schedules are required to ensure that when ears are harvested, the hybrids achieve maturity by accumulating enough growing degree units (GDU), a temperature index used to estimate plant progression, and there is a consistent weekly harvest quantity which is below the storage capacity. Poor scheduling may result in having harvest quantities above the maximum capacity or cause growers to experience logistical challenges due to inconsistent harvest quantities.

Many factors affect the planting date of crops such as weather, soil temperature, and planting resources. Traditionally, farmers and growers schedule their planting dates in a way that they will have a continuous harvest of crops at the end of growing season [3]. Such a planing schedule has multiple benefits: (1) farmers can better schedule their harvest dates without having to harvest all crops at a time, and (2) the price of crops may change depending on the demand and farmers can decide when to harvest to maximize their profit. Farmers usually extend the harvesting period by either planting varieties with different days to maturity or planting a variety multiple times in succession. For example, farmers wait until first planted sweet corns are about 1 to 2 inches tall before planting the next batch [3]. However, these methods are not optimal because they do not consider important factors such as accumulated GDU, planting window, and storage capacity.

In this paper, we propose two mixed integer linear programming (MILP) models which provide optimal planting and harvest schedules while ensuring consistent weekly harvest quantities that are below the maximum storage capacity. Our proposed optimization method considers the uncertainty in the GDUs and provides optimal planting and harvest schedules under different GDU possibilities.

More recently, deep learning techniques have been utilized in many agricultural big data applications including crop yield prediction, classification of crop tolerance to heat and drought, and image-based crop yield estimation [4, 5, 6, 7, 8, 9, 10, 11]. There are very few studies in the literature that use optimization models to obtain optimal planting or harvest schedule. In crop planning, Cid-Garcia et al. and Sarker et al. used linear programming model to help farmers decide how to dedicate different parts of their land to different crops at different points of time to maximize their profit [12, 13]. To the best of our knowledge our paper is the first study to propose a MILP model which optimizes both planting and harvesting dates.

## 2 Problem Statement

In the current century, scientists are confronted with the grand challenge of increasing agricultural production to secure the world’s food supply and feed the growing population. There are two broad options that can address the aforementioned global challenge: (1) increasing the farm allocation areas or (2) increasing crop production on existing agricultural land [14]. Due to the fact that bringing new land into production can cause large-scale disruption of existing ecosystems associated with them and increase greenhouse gas emissions, improving productivity on the existing farmland is desirable. Commercial hybrids can be named as one the promising ways that can increase crop production.

The development and widespread use of commercial hybrids preceded a rapid rise in the corn yields. This dramatic yield improvement can cause over capacity problems for storage facilities. Moreover, different corn hybrids have different planting windows and poor planting and harvest schedules may result in having inconsistent weekly harvest quantities. An erratic weekly harvest quantity is not preferable as it makes it difficult to organize the logistics of transporting the crop and create productivity issues. In order to ensure that the number of harvested ears in each week does not exceed the storage capacity, and fallows a consistent pattern, an optimization model is essential for striking a balance between the storage capacity and a minimum number of harvesting weeks. To this end, an optimization model is required to find the optimal planting and harvest schedules which minimizes the difference between the weekly harvest quantity and the storage capacity while guarantees a consistent pattern of weekly harvest.

In the 2021 Syngenta Crop Challenge [15], Syngenta provided participants with real-world data including historical daily GDU of two sites namely site 0 and site 1, planting windows, required GDUs, and harvest quantities of 1375 and 1194 corn hybrids planted in site 0 and site 1 respectively. Then, participants were asked to use this data to determine optimal planting and harvest schedules of these corn hybrids under two storage capacity scenarios so that we are not over holding capacity in harvesting weeks and there is a consistent harvest quantity each week. For scenario 1, a maximum storage capacity of 7000 ears and 6000 ears are defined for site 0 and site 1 respectively. For scenario 2, there is not a predefined capacity and the optimization model should determine the lowest storage capacity. For both scenarios, the optimized planting and harvest schedules should result in achieving a minimum number of harvesting weeks and consistent weekly harvest quantities which are below the storage capacity.

Growing Degree Days (GDD), also known as Growing Degree Units (GDU) or heat units are a measure of accumulation of heat or temperature units used to estimate the plant growth stage and are calculated based on air temperature by subtracting the base temperature from the average of the daily maximum and minimum air temperatures in °F or °C. Base temperature is the temperature below which the crop does not grow and it is different for different types of crops. In the case of corn, the base temperature is 50°F (10°C), and the equation for calculating GDU in °F is [16]:

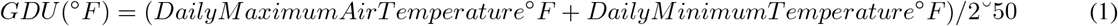

If the daily maximum temperature is above 86°F (30°C), then the daily maximum temperature is set at 86°F (30°C) as above that temperature the growth rate of corn is not significant. Likewise, when the daily minimum temperature is less than 50°F (10°C), then this value is set at 50°F (10°C) [17].

In this paper, we focus on the planting and harvest schedules of two separate groups of corn hybrids each including different corn seed populations planted in two sites (site 0 and site 1). Our optimization models aim to (1) obtain optimal planting and harvest schedules for each seed population planted in site 0 and site 1 to ensure consistent weekly harvest quantities that don’t exceed the maximum capacity of each site, and (2) identify the lowest capacity required for site 0 and site 1 in which the optimal planting and harvest schedules for each seed population give us consistent weekly harvest quantities during minimum number of harvesting weeks.

## 3 Data

In the 2021 Syngenta Crop Challenge, participants were asked to use real-world data to determine optimal planting and harvest schedules of two separate groups of corn hybrids planted in two sites. The dataset included information of 1375 and 1194 different corn seed populations planted in site 0 and site 1 respectively.

Harvest quantities of each of these seed populations were provided for each capacity scenario and their distributions are shown in figures 1 and 2 for site 0 and site 1 respectively. Figures 1 and 2 show that most harvest quantities for scenario 1 are close to 250, and for scenario 2 are close to 400 for both sites. As it its shown in these figures the spread of harvest quantities for scenario 1 and scenario 2 is notably different in both sites. The harvest quantities for scenario 1 mostly fall between 45 – 650 and between 60 – 600 for site 0 and site 1 respectively while for scenario 2 those ranges are 40 – 850 and 60 – 820 for site 0 and site 1 respectively. In short, these histograms show that higher values are more common for scenario 2 in both sites.

**Figure 1:**
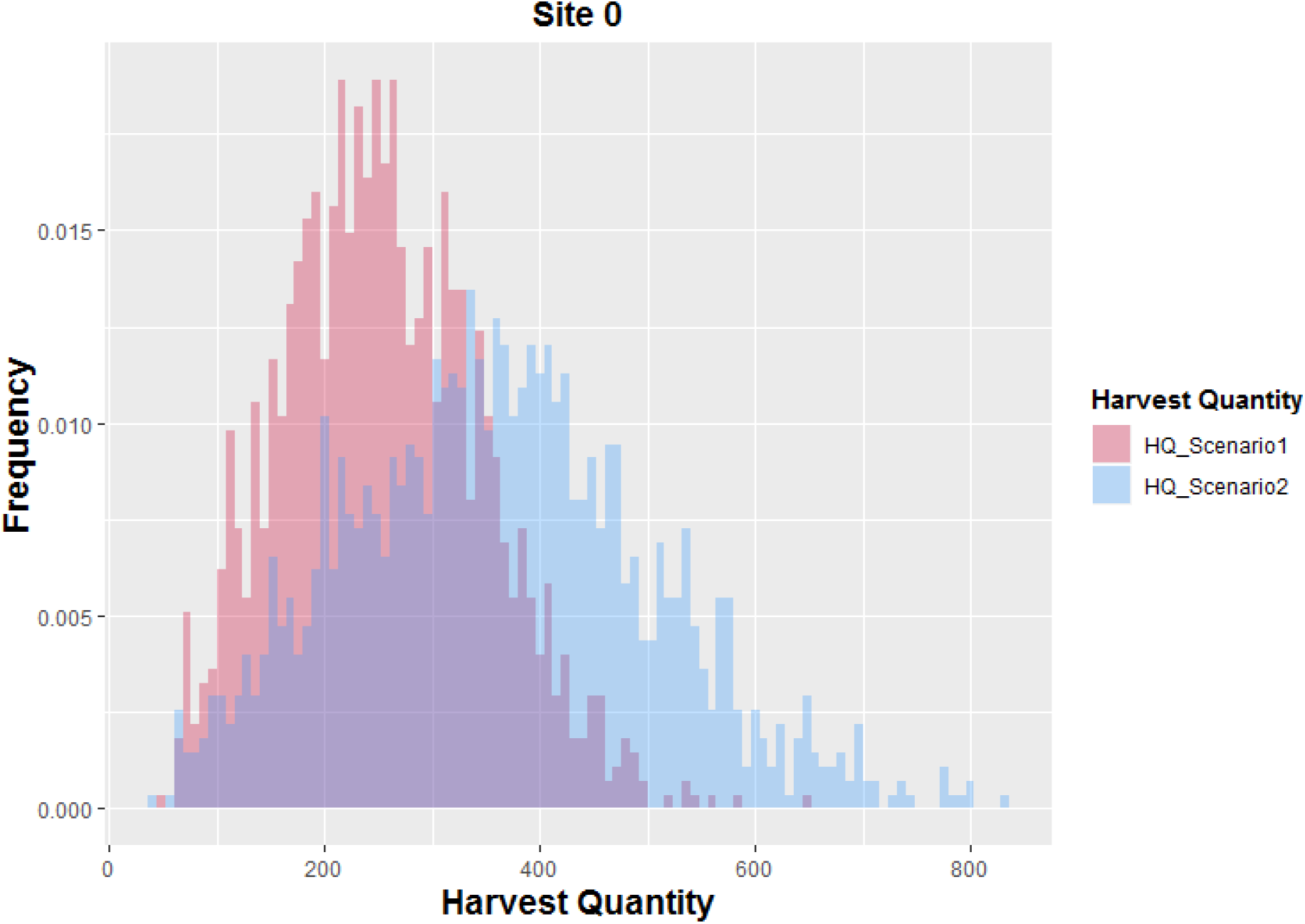
Harvest quantity distributions of 1375 seed populations planted in site 0 for scenario 1 and scenario 2.

**Figure 2:**
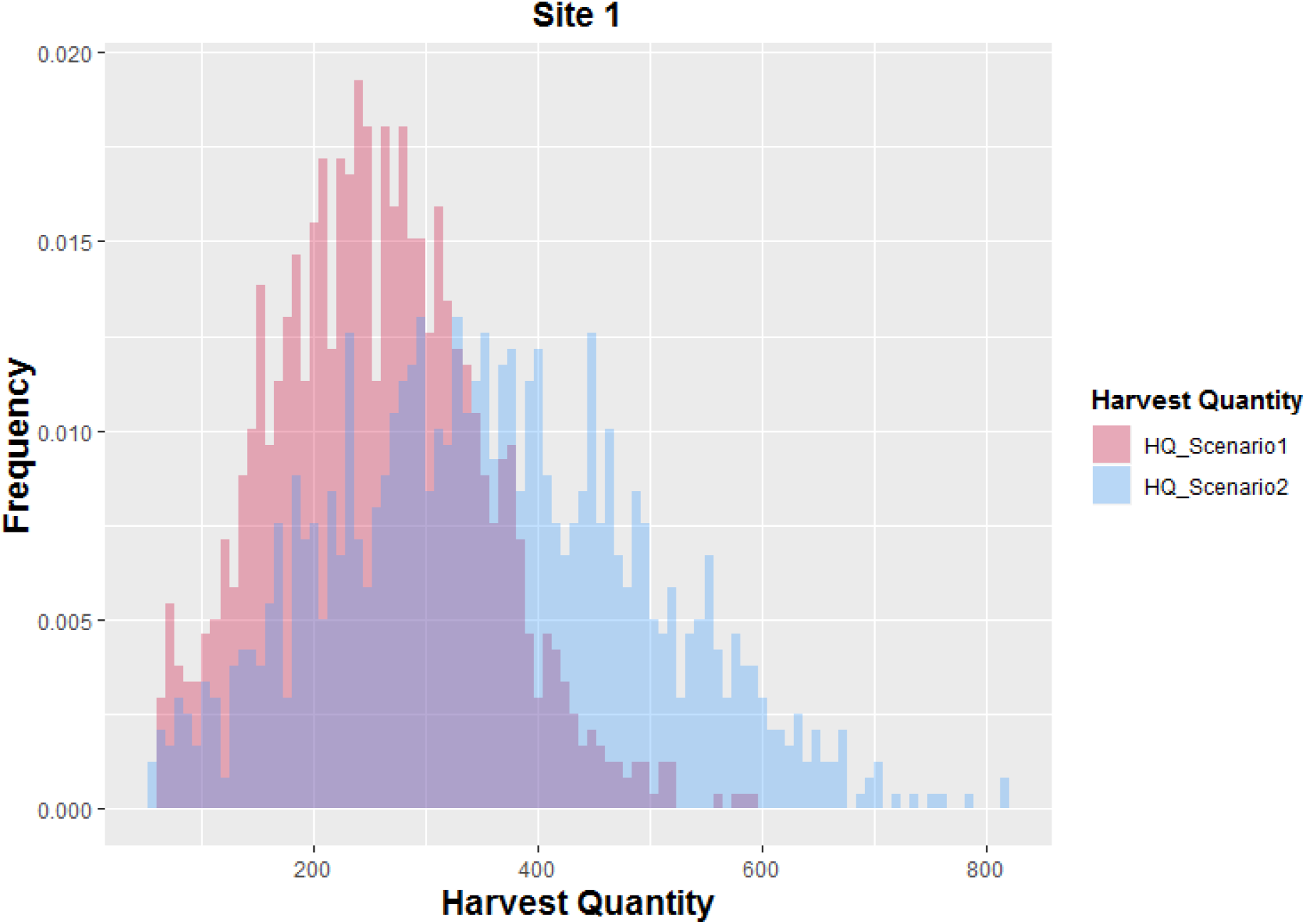
Harvest quantity distributions of 1194 seed populations planted in site 1 for scenario 1 and scenario 2.

Due to the fact that the planting date is one of many yield influencing factors [18] and the right planting window can help to achieve the highest potential yield [19] the earliest and latest planting dates of each seed population were given.

The growing degree units in Celsius were provided for each site for each day over the last 10 years from 2009 to 2019. Also, the dataset contained the required GDU for each seed population planted in site 0 and site 1. Required GDU is a certain number of growing degree units which is required in order for each corn population to reach maturity and be ready for harvesting.

## 4 Method

### 4.1 Data Preprocessing

As it was discussed in section 3 the early and late planting dates for each seed population planted in each site were provided. Daily data is difficult to work with because it makes our optimisation models hard to solve as it increases the size of the model. Also, in our optimization models we used cumulative weekly GDUs, so the planting windows needed to be consistent with them. To avoid these problems, we converted the early and late planting dates to the week number corresponding to each date. In order to convert the dates to week number corresponding to them we used Microsoft Excel WEEKNUM function in which week 1 begins on January 1, and the next weeks begin on Sundays. The reason of using this function is that in the 2021 Syngenta crop challenge it was assumed that each week runs from Sunday – Saturday.

Figures 3 and 4 show the weekly planting window of each of 1375 and 1194 populations planted in site 0 and site 1 respectively which are used for both scenarios in our optimization models.

**Figure 3:**
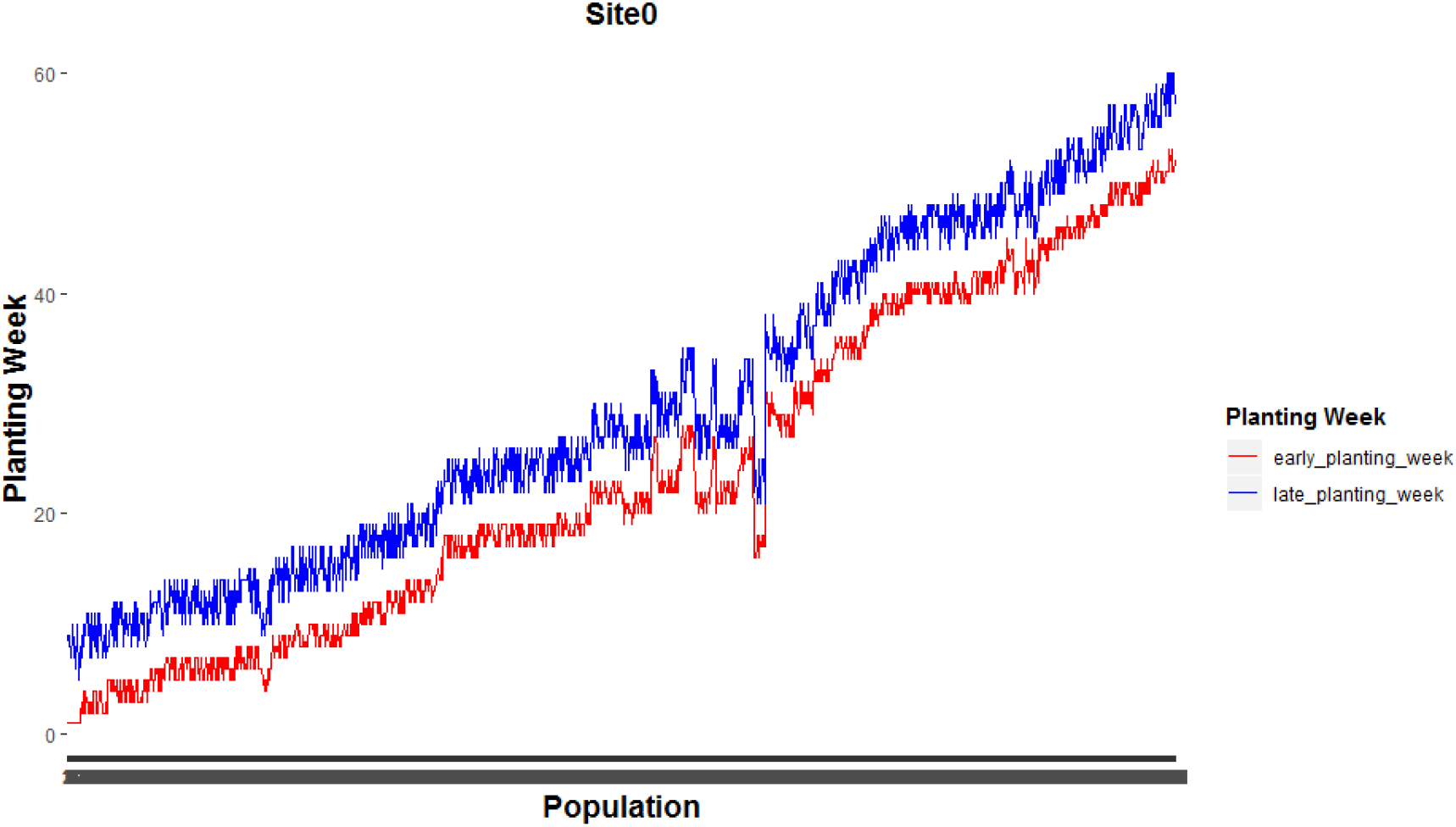
Weekly planting windows of 1375 seed populations planted in site 0.

**Figure 4:**
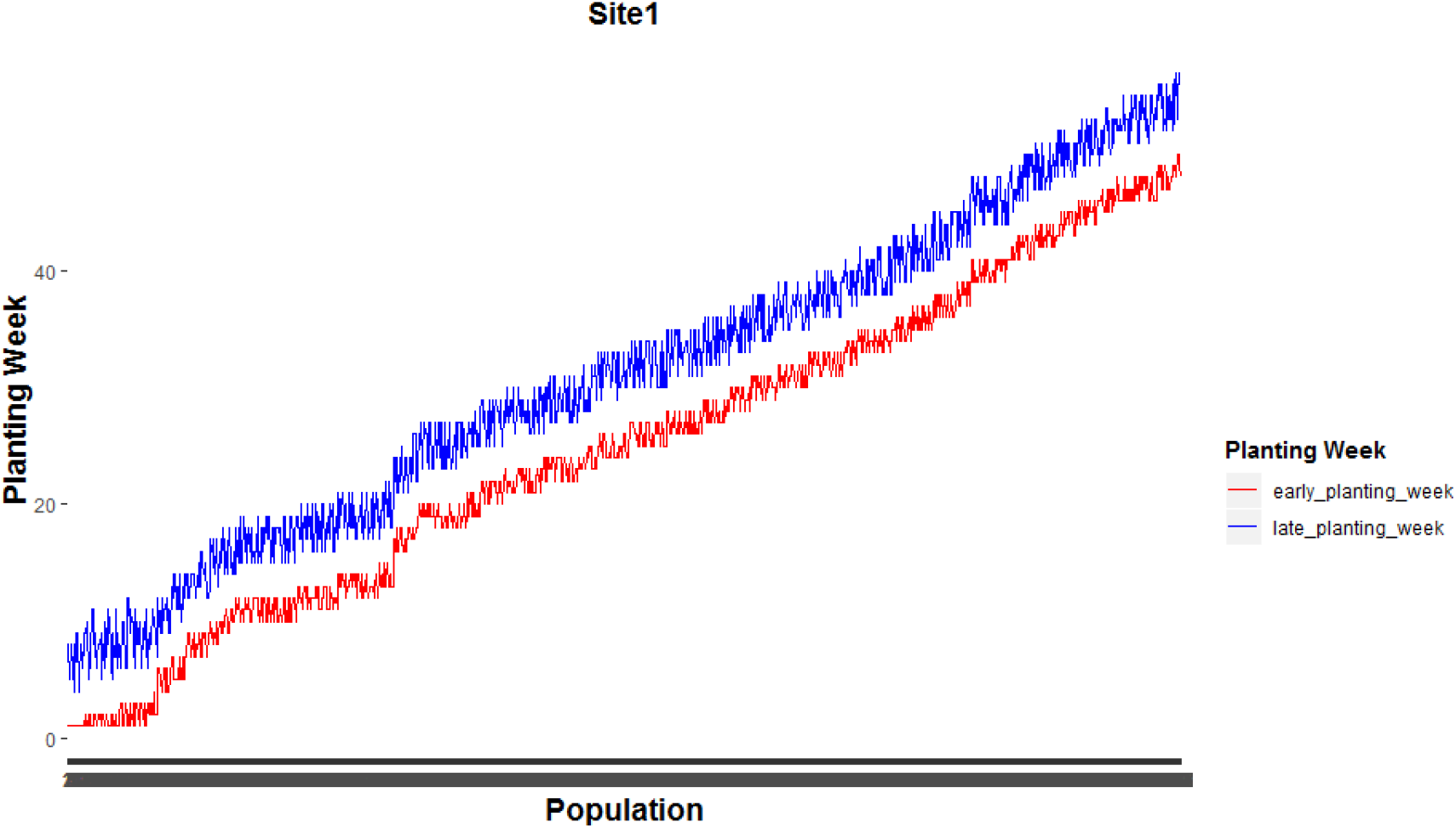
Weekly planting week windows of 1194 seed populations planted in site 1.

### 4.2 GDU Possibilities

GDU estimation is an inevitable part of scheduling the planting and harvesting dates to have consistent harvest quantities during limited number of weeks, because each seed population should accumulate a certain number of GDUs to be ready to be harvested but it is unknown a priori. In this section, we describe our approach for the GDU estimation for site 0 and site 1 and apply it to estimate the weekly GDUs during 70 weeks after January 1st 2020 using the given historical daily growing degree units from 2009 to 2019 for each site.

Based on equation 1 temperature is the most important dominating factor affecting the GDU and the larger GDU indicates the higher average temperature and the larger amount of heat available for the growth of crops. The boxplots of the average weekly GDU of site 0 and site 1 during the last 10 years from 2009 to 2019 are shown in Figure 5. This figure shows that the average weekly GDU varies with sites and site 0 is located in a place with lower average temperature as the median average weekly GDU of site 0 is less than the lower quartile of the average weekly GDU of site 1 (that is, over three-quarters of the average weekly GDU of site 1 are greater than the median average weekly GDU of site 0). The mean of the average weekly GDUs (shown by the white circle in each boxplot in figure 5) of site 0 is also less than that of site 1.

**Figure 5:**
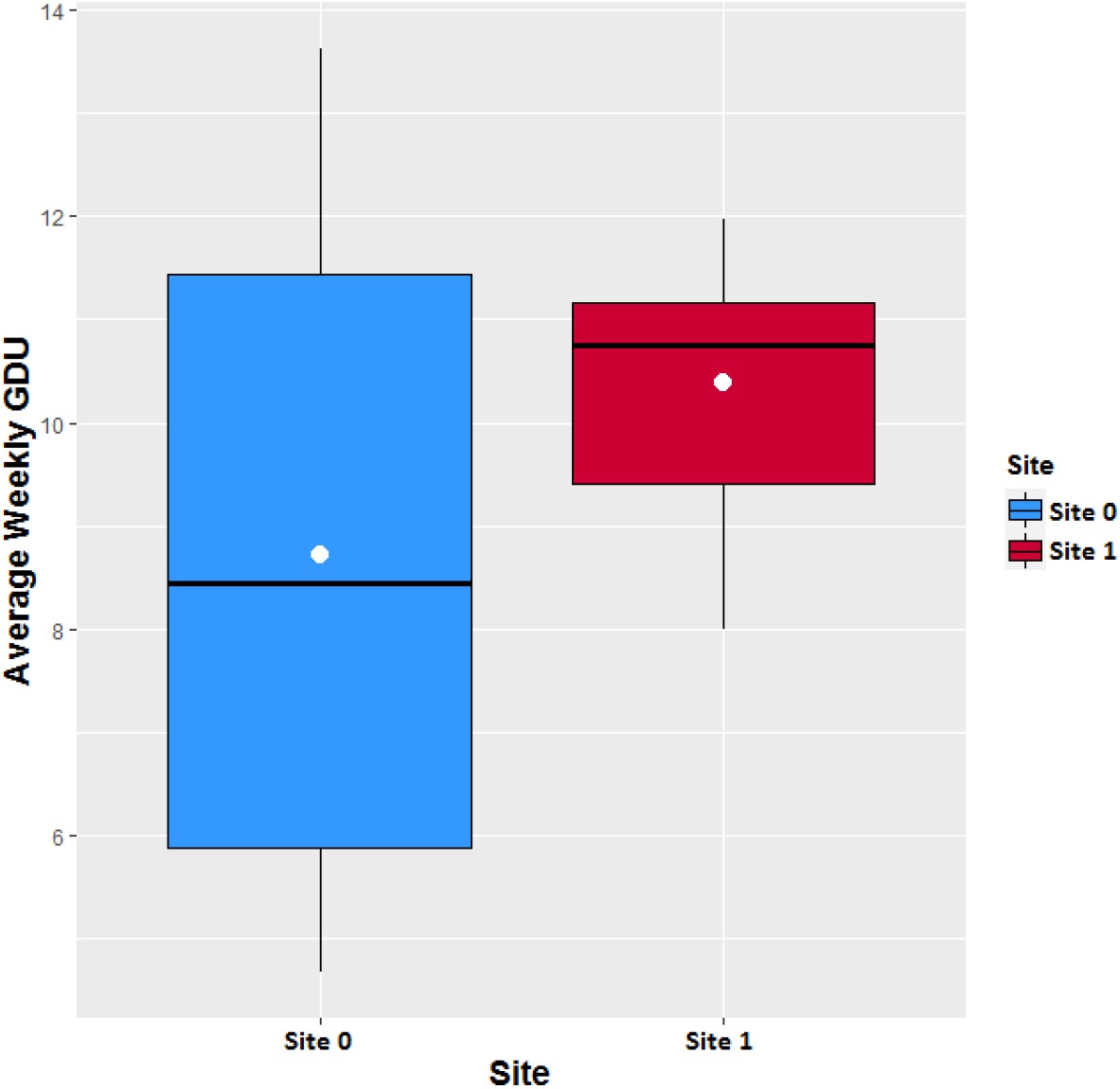
Box plot of the average weekly GDU during the last 10 years from 2009 to 2019 of each site. The white circle in each boxplot is the mean of the average weekly GDU during the last 10 years from 2009 to 2019.

Because of the lower amount of heat available for the growth of crops in site 0 the growing degree units required in order for the corn population to achieve maturity accumulate slower and the seed populations may need more than 70 weeks to be ready to be harvested. So, for site 0 first we found the minimum weekly GDUs for each of 70 weeks that enable us to harvest the whole 1375 seed populations planted in site 0 in 70 weeks. The minimum GDUs for the 70 weeks are found using the daily historical GDUs from 2009 to 2019. For this purpose, first of all we calculated the sum of the GDUs during all days of each week (weekly GDUs) for each year from 2009 to 2019. For site 0, 80 percentile values of the weekly GDUs over the last 10 years of each of 52 weeks were found the minimum weekly GDUs for the next 52 weeks. Note that the same values of week 1 to week 18 were used for weekly GDUs for week 53 to week 70. We utilized 85 percentile, 90 percentile, and maximum values in case of having higher average temperature in the planning year for site 0. On the other hand, because site 1 has a larger amount of historical weekly GDUs, the minimum weekly GDUs for the 52 weeks that enable us to harvest the whole 1194 seed populations planted in site 1 in 70 weeks was found to be the 5 percentile value of the weekly GDUs over the last 10 years (The same values of weekly GDUs from week 1 to week 18 were used for week 53 to week 70). We used first quartile, median, and third quartile in case of having lower or higher averge temperature in a planning year for site 1. Succinctly, in order to take different weather conditions (average temperatures) in a planning year into account, we used three GDU possibilities as estimations of the weekly GDUs including 80 percentile, 90 percentile, and maximum of the past 10 years weekly GDUs for site 0 and first quartile, median, and third quartile of the past 10 years weekly GDUs for site 1.

### 4.3 Proposed Optimization Models

We propose two optimization models to schedule planting and harvesting dates of each seed population planted in site 0 and site 1 under the following scenarios: (1) the planting and harvesting dates of each seed population planted in each site should be scheduled in a way that the weekly number of harvested ears during 70 weeks after January 1st 2020 does not exceed the capacity of site 0 and site 1 which are 7000 and 6000 ears respectively. (2) there is not a predefined capacity, and we should determine planting and harvesting dates of each seed population and the lowest capacity required for each site to reach consistent weekly harvest quantities which are below the capacity during 70 weeks after January 1st 2020.

As it was explained in section 3, 1375 and 1194 seed populations planted in site 0 and site 1 respectively. The harvest quantities of each of these seed populations planted in each site varies with scenarios. Other given data including historical daily GDUs of each site, planting window and the required GDUs of each seed population are the same in both scenarios.

#### 4.3.1 Optimization Model for Scenario 1

This subsection presents our proposed optimization model for corn planting and harvest scheduling for scenario 1 which results in consistent weekly harvest quantities that are below the maximum capacity. Our optimization model consists of the following decision variables:

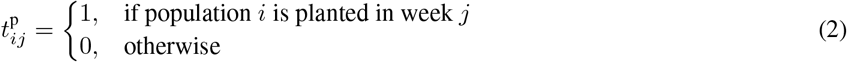

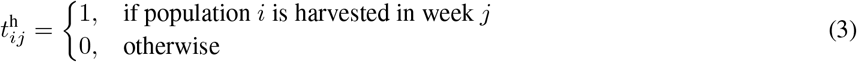

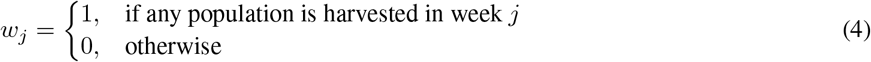

The notations used in our optimization model include:

- *C*: the storage capacity
- *GDU_j_*: the cumulative weekly GDU which are accumulated till week *j*
- *N*: the total number of populations
- *T*: number of weeks after Jan 1st of planning year (it is 70 weeks for the 2021 Syngenta crop challenge.)
- *HQ_i_*: number of ears (harvest quantity) produced by population *i*
- 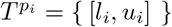: the planting window for the population *i*, where *l_i_* and *u_i_* are the corresponding earliest and latest planting dates, respectively.
- 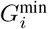: number of growing degree units needed before harvesting population *i* (required GDUs for population *i*)
- *θ_w_*: the coefficient for the number of weeks used for harvesting
- *M*: constant large number

As such, we define our optimization model as following:

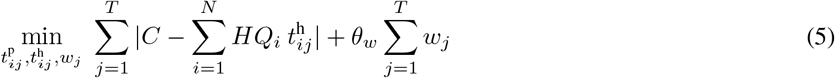

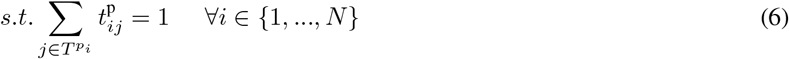

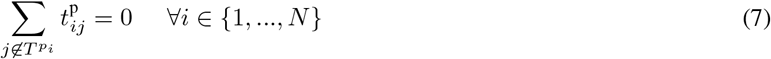

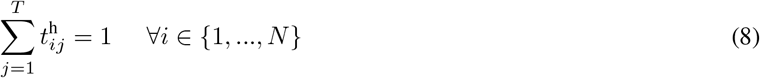

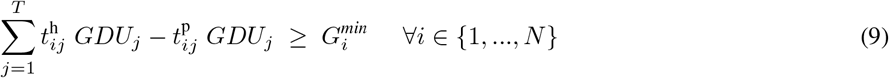

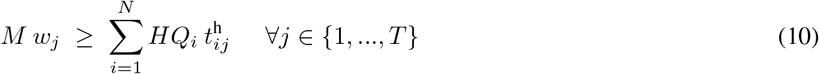

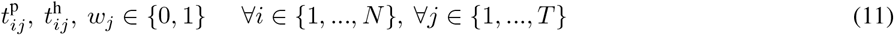

Here, the objective (5) is to minimize the difference between the weekly harvest quantity and the capacity for each harvesting week while using the minimum number of weeks for harvesting. We adopt the absolute value function in our objective function because it can be easily linearized and is computationally more tractable [20]. Constraints (6) and (7) make sure that each population is planted within its corresponding planting window. Constraint (8) means that each population can only be harvested in one week. Constraint (9) enforces the model to harvest populations only if they have accumulated the least amount of required GDU. Constraint (10) ensures “only if” direction: if any population is harvested in week *j*, then *w_j_* = 1. Finally, Constraint (11) indicates the appropriate types of the decision variables.

Due to having the absolute value function inside our objective, the above-mentioned model is a nonlinear optimization problem which is hard to solve. As a result, we reformulate our model into an equivalent mixed integer linear program (MILP) by introducing a set of new variables which is as follows [20]:

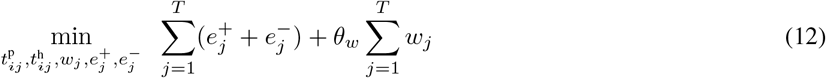

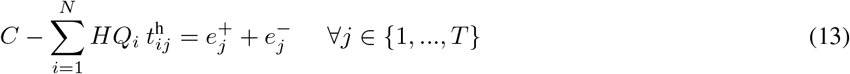

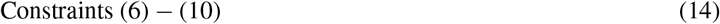

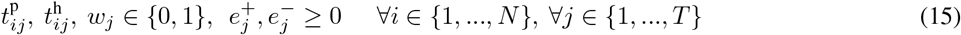

Where, the objective (12) is to minimize the positive and negative errors between the storage capacity and the sum of harvested quantities for each week which is equivalent to objective (5). Constraint (13) defines the two error terms where at least one of them should be zero in the optimal solution. Constraint (15) indicates the appropriate types of the decision variables.

#### 4.3.2 Optimization Model for Scenario 2

This subsection presents our proposed optimization model for scenario 2 where there is not a predefined capacity, and the goal is to determine planting and harvesting dates of each population (during *T* weeks) and also the lowest capacity required for the whole populations planted in each site. This optimization model has the same decision variables and constraints with the optimization model proposed for scenario 1, but the objective function is changed to achieve the goal of scenario 2. The optimization model for scenario 2 is as follows:

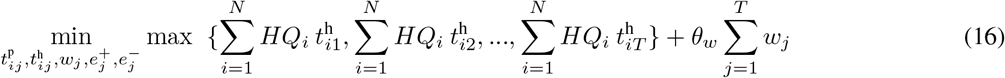

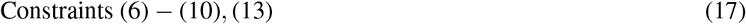

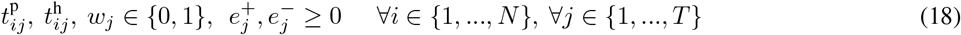

This is a Minimax Linear Programming Problem (MLPP), and the objective (16) is to minimize the maximum amount of weekly harvest which simultaneously minimize the amount of weekly harvests for all weeks while using the minimum number of weeks for harvesting. Since the objective (16) is nonlinear, we reformulate our model into an equivalent mixed integer linear program (MILP) by introducing a new variable denoted by *z* [21]:

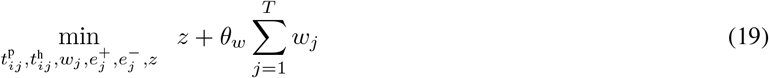

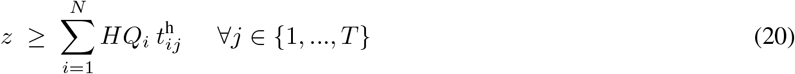

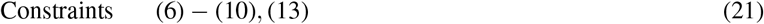

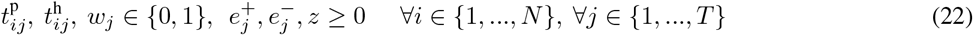

Here, the objective is to minimize the summation of the maximum value of the weekly harvest quantities and the total number of harvesting weeks. Constraint (20) ensures that the maximum value of the weekly harvest quantities is always greater than or equal to amount of harvests of each week.

## 5 Results

In this section, we present the quantitative results of our optimization models for scenario 1 and scenario 2 under three GDU possibilities. In the 2021 Syngenta crop challenge, participants were asked to schedule the planting date of each seed population sometimes within the given planting window for each seed population and the harvesting dates during 70 weeks after January 1st 2020. As it was discussed in section 4.2 the daily GDUs of these 70 weeks were unknown a priori and as a part of the challenge we made use of historical daily GDUs from 2009 to 2019 provided for each site to estimate GDUs of these 70 weeks for each site. In order to consider different weather conditions in a planning year, we solved each optimization model under 3 GDU possibilities including 80 percentile, 90 percentile, and maximum of the previous weekly GDUs from 2009 to 2019 for site 0, and first quartile, median, and third quartile of the previous weekly GDUs from 2009 to 2019 for site 1. We could not use 1st, 2nd, or 3rd quartile of the previous weekly GDUs for site 0 because the low GDUs values resulted in the infeasibility of the model due to the GDU constraint (9).

We implemented our MILP models (12)-(15) and (19)-(22) in MATLAB R2018a and solved with MILP commercial solver Gurobi Optimizer.

### 5.1 Results of the Optimization Model for Scenario 1

The MILP model of scenario 1 (12)-(15) was run 6 times for each site under 3 GDU possibilities with their own input variables. Input variables for the optimization model for each site include: the storage capacity (*C*) of 7000 ears for site 0 and 6000 ears for site 1, total number of populations (*T*) of 1375 planted in site 0 and 1194 planted in site 1, planting window (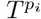) including the early and late planting week for each seed population planted in site 0 and site 1, the number of ears produced by each population (*HQ_i_*) planted in site 0 and site 1 given for scenario 1, and required GDUs of each seed population 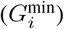 planted in site 0 and site 1. Because the sum of 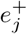 *and* 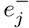 in equation (12) is the absolute value of the difference between the weekly harvest quantity and the capacity for each harvesting week, it can take value up to 7000 and 6000 for weeks with no harvest quantities for site 0 and site 1 respectively. Besides, *w_j_* is a binary variable. As a result, to avoid ignoring minimizing the number of weeks by the model, we used a coefficient for the number of weeks used for harvesting (*θ_w_*) and assign it to 7000 for site 0 and 6000 for site 1. The cumulative weekly GDU (*GDU_j_*) of each site are calculated using the estimated weekly GDU based on the 3 GDU possibilities for each site. The constant large number (*M*) and number of weeks after Jan 1st of planning year (*T*) are equal to 20000 and 70 respectively for both sites.

Weekly harvest quantities under 3 GDU possibilities are shown in figures 6 and 7 for site 0 and site 1 respectively, which suggest that the proposed MILP model (12)-(15) was able to schedule the planting and harvesting dates of the whole 1375 and 1194 populations planted in site 0 and site 1 respectively with different planting window, required GDUs, and harvest quantities in a way that resulted in consistent weekly harvest quantities that are below the storage capacities.

**Figure 6:**
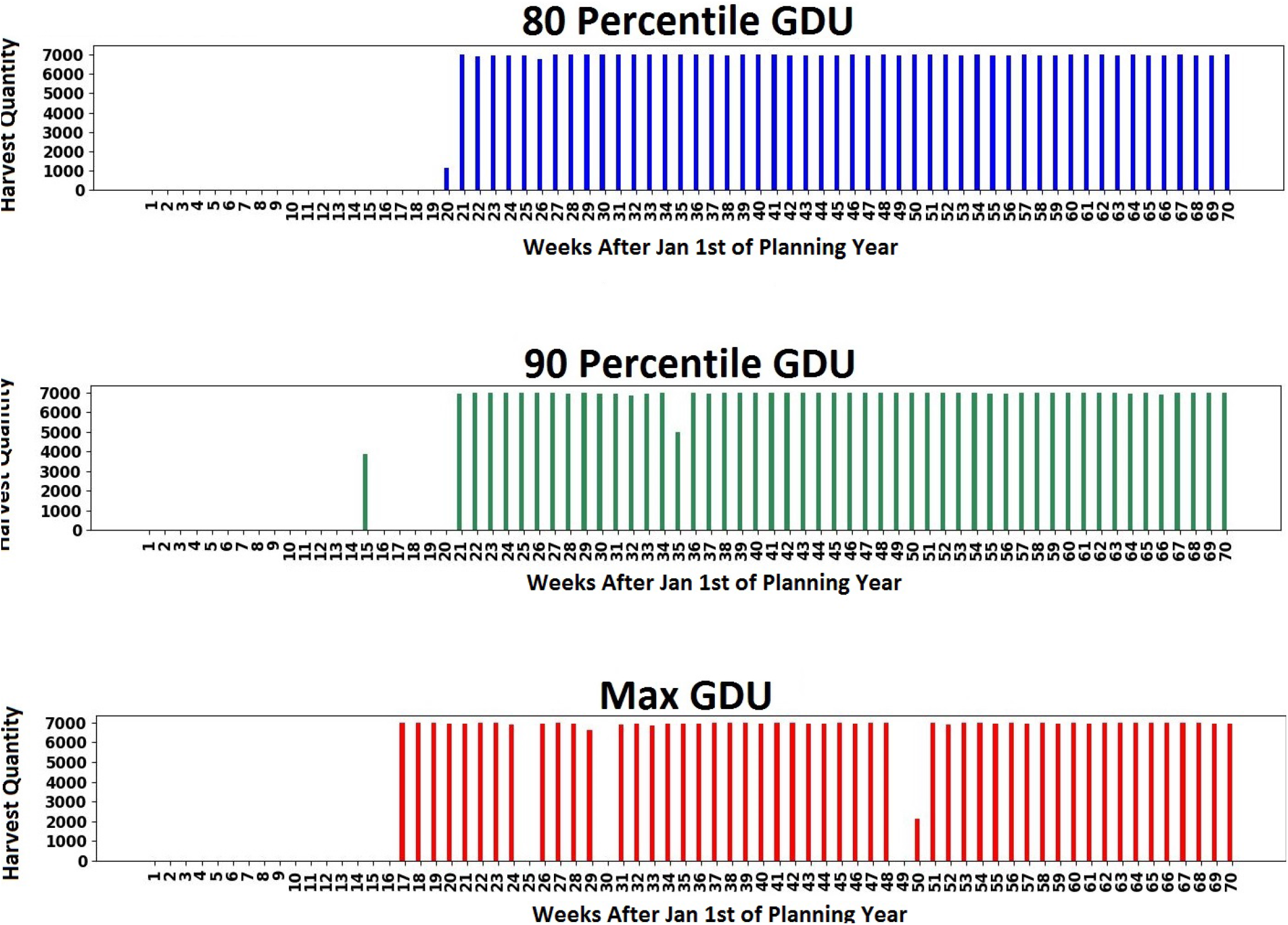
Weekly harvest quantity of 1375 seed populations planted in site 0 with a capacity of 7000 ears for scenario 1 under 3 GDU possibilities.

**Figure 7:**
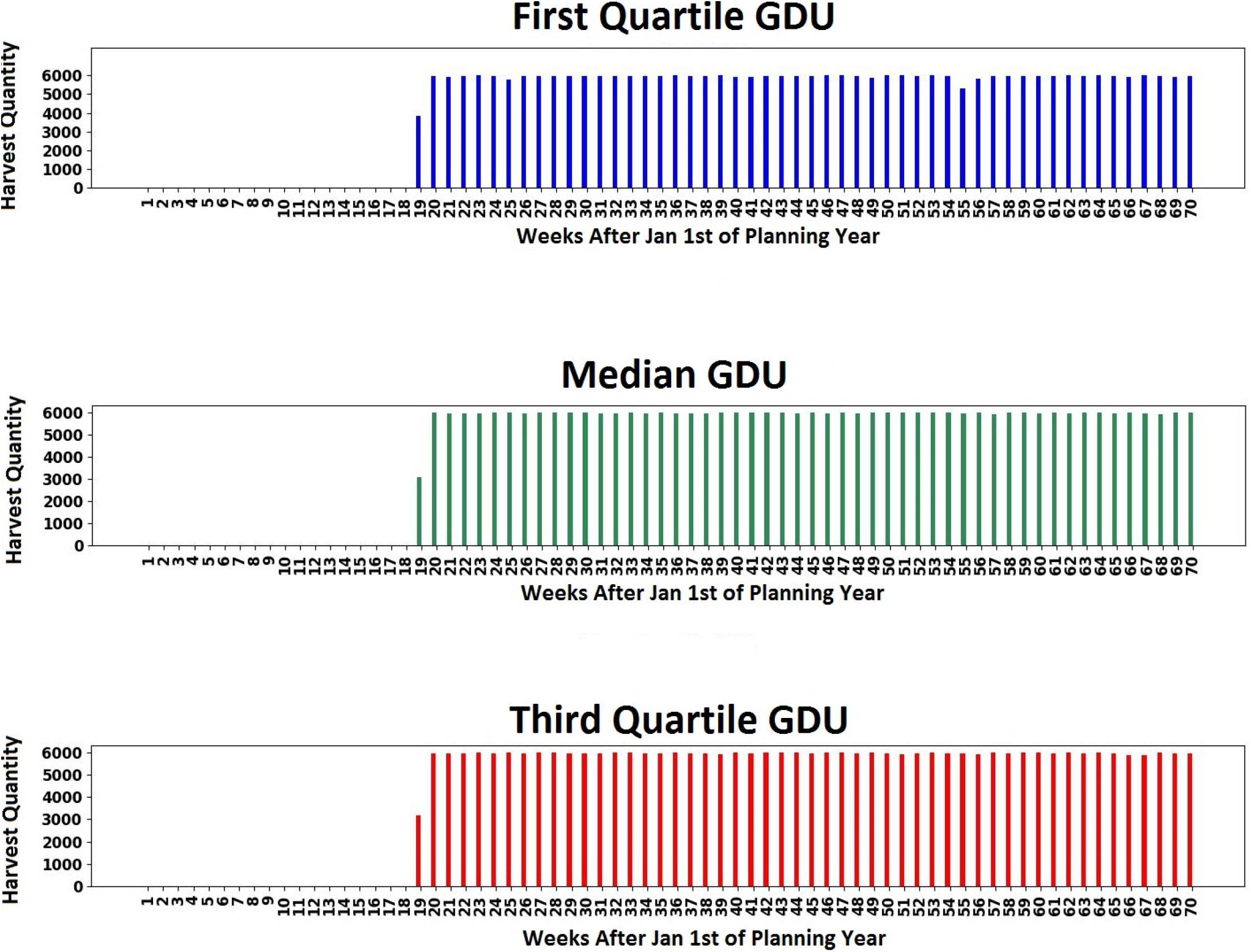
Weekly harvest quantity of 1194 seed populations planted in site 1 with a capacity of 6000 ears for scenario 1 under 3 GDU possibilities.

As it is shown in figures 6 and 7 GDU possibilities do not affect the total number of harvesting weeks and it is equal to 51 weeks and 52 weeks for all 3 GDU possibilities of site 0 and site 1 respectively.

Because the total number of harvesting weeks is equal for 3 GDU possibilities of each site, we extracted the weeks without harvest quantities and plotted the boxplots of the *log*10 of difference between the weekly harvest quantity and the capacity of each site (capacity of 7000 ears of site 0 and 6000 ears of site 1) for all harvesting weeks for 3 GDU possibilities. These boxplots shown in figures 8 and **??**help us to see how the absolute maximum and absolute median difference between the weekly harvest quantity and the capacity change using different GDU possibilities. The absolute maximum and absolute median difference between the weekly harvest quantity and the capacity among all harvesting weeks are shown in table 1 for each GDU scenario for site 0 and site 1. Table 1 indicates that 90 percentile and 80 percentile GDU possibilities for site 0 have the minimum absolute maximum and absolute median difference between the weekly harvest quantity and the capacity respectively. Among GDU possibilities of site 1, first quartile has the the minimum absolute maximum and absolute median difference between the weekly harvest quantity and the capacity.

**Table 1:**
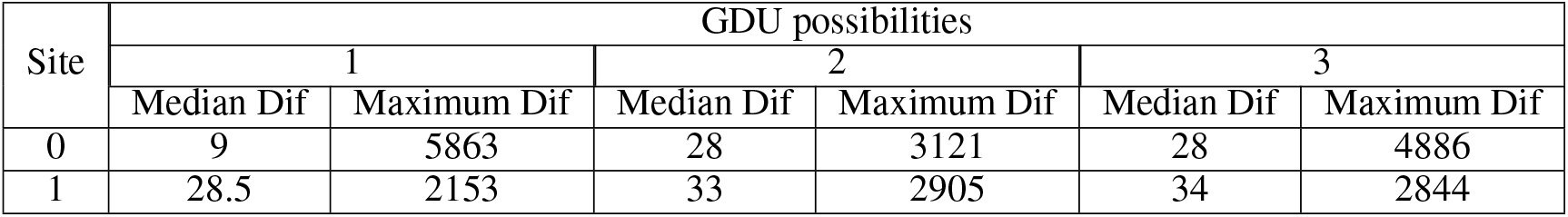
The absolute median difference between the weekly harvest quantity and the capacity (Median Dif) and the absolute maximum difference between the weekly harvest quantity and the capacity (Maximum Dif) among all harvesting weeks for site 0 and site 1 under 3 GDU possibilities. For site 0 GDU possibilities from 1 to 3 are: 80 percentile, 90 percentile, and maximum respectively. For site 1 GDU possibilities from 1 to 3 are: first quartile, median, and third quartile respectively.

**Figure 8:**
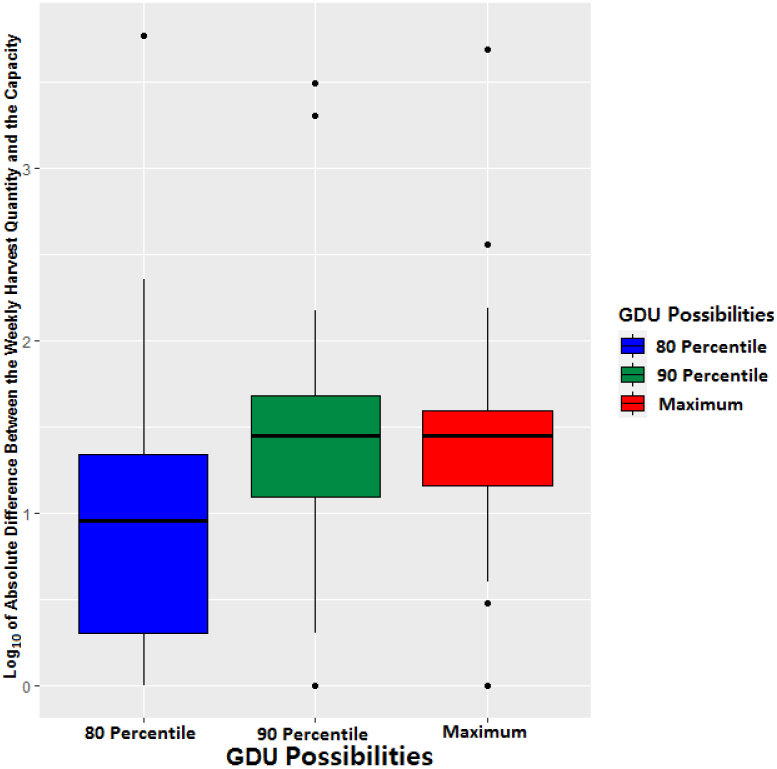
Box plot of log of absolute difference between the weekly harvest quantity and the capacity of 7000 ears among all harvesting weeks for 3 GDU possibilities of site 0.

**Figure 9:**
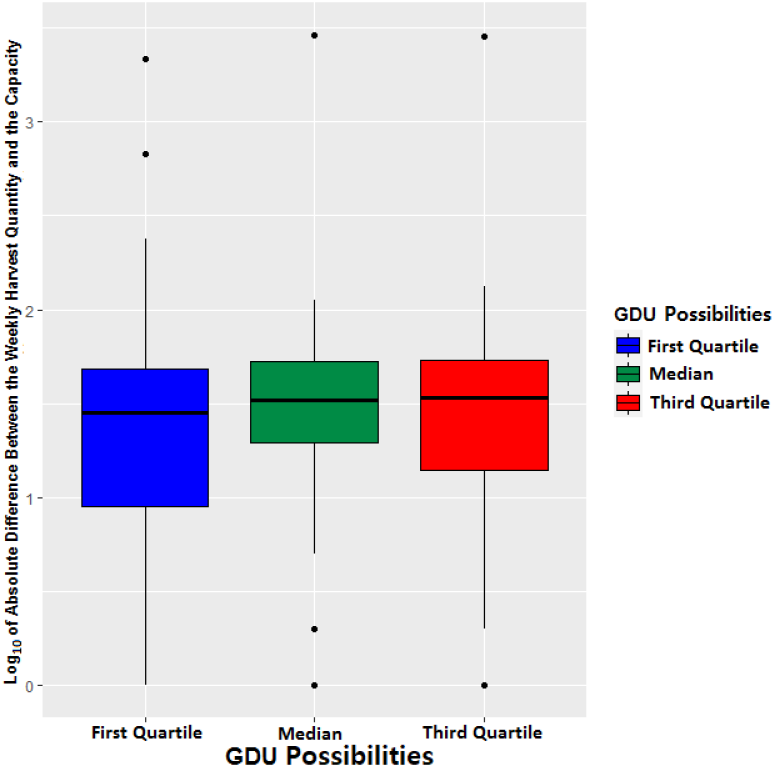
Box plot of log of absolute difference between the weekly harvest quantity and the capacity of 6000 ears among all harvesting weeks for 3 GDU possibilities of site 1.

Figures 10 and 11 present optimal planting and harvesting weeks suggested by our proposed MILP model (12)-(15) for a subset from the whole 1375 seed populations planted in site 0 and 1194 seed populations planted in site 1 respectively for each of 3 GDU possibilities.

**Figure 10:**
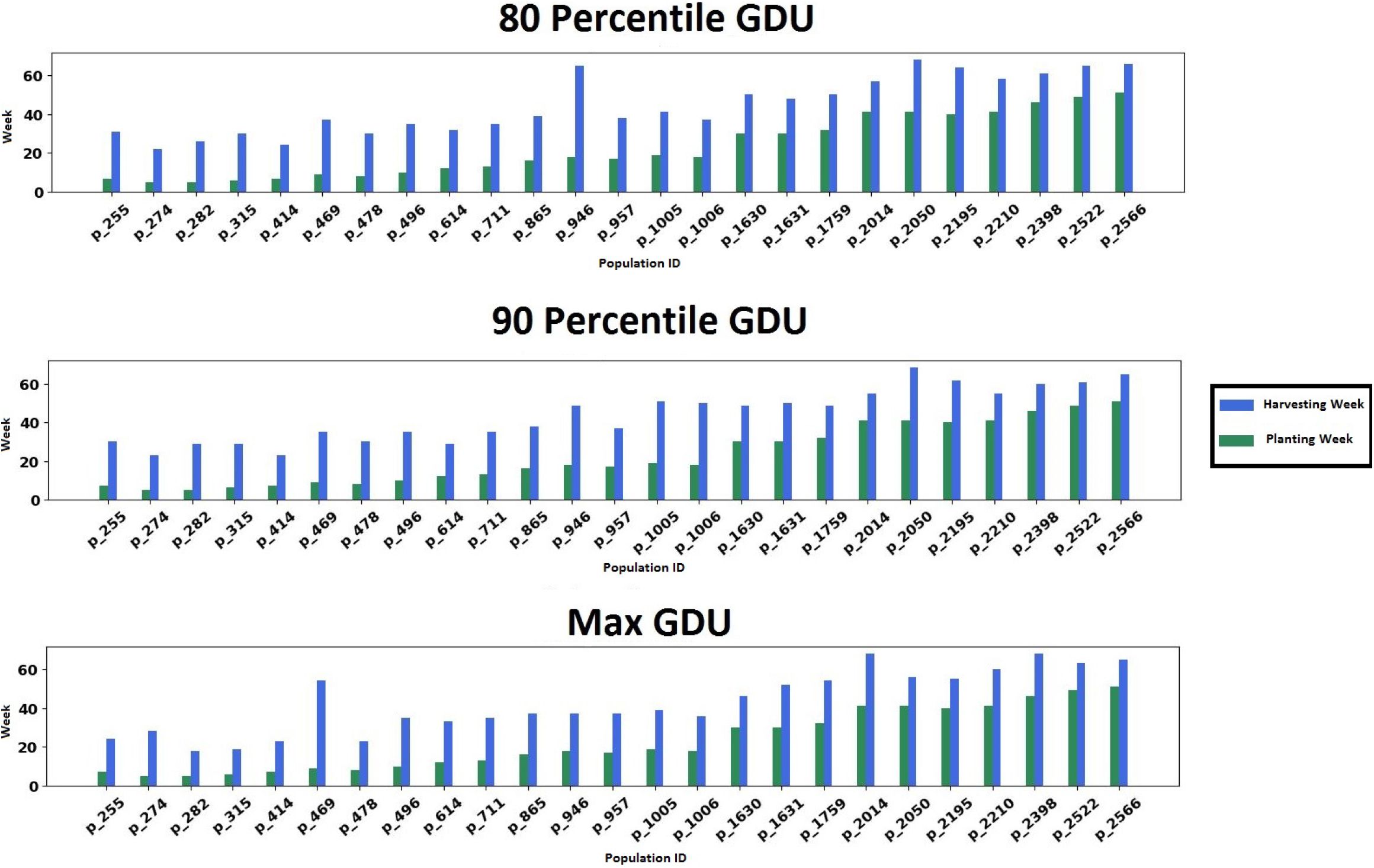
Optimal planting and harvesting week for a subset from the whole 1375 seed populations planted in site 0 under 3 GDU possibilities for scenario 1.

**Figure 11:**
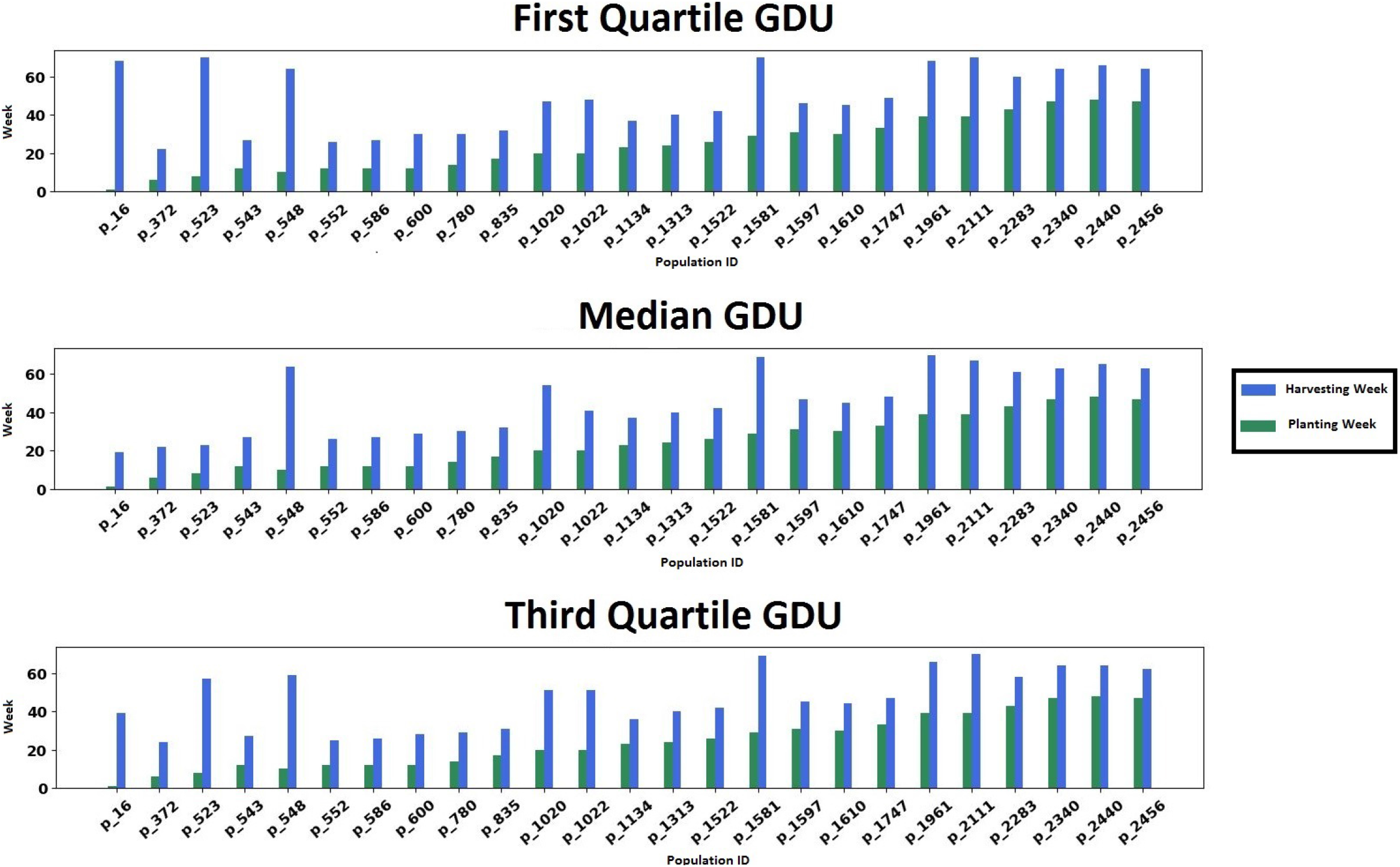
Optimal planting and harvesting week for a subset from the whole 1194 seed populations planted in site 1 under 3 GDU possibilities for scenario 1.

### 5.2 Results of the Optimization Model for Scenario 2

The same input variables as the MILP model of scenario 1 (12)-(15) were used in the MILP model of scenario 2 (19)-(22) except the number of ears produced by each population (*HQ_i_*) planted in site 0 and site 1, and the coefficient for the number of weeks used for harvesting (*θ_w_*). For this scenario in which the model is supposed to determine the lowest capacity required for each site, we are given different quantities of the number of ears produced by each population (*HQ_i_*) planted in site 0 and site 1. Moreover, here we set (*θ_w_*) to be 1. Because of the complexity of this MILP model (19)-(22) we used stopping criterion of 0.1 % optimality gap for each site and scenario.

The suggested lowest capacity required for each site under each GDU scenario by the MILP model (19)-(22) is presented in table 2.

**Table 2:** The the lowest capacity required for site 0 and site 1 under 3 GDU possibilities. For site 0 GDU possibilities from 1 to 3 are: 80 percentile, 90 percentile, and maximum respectively. For site 1 GDU possibilities from 1 to 3 are: first quartile, median, and third quartile respectively.

| Site | GDU possibilities |  |  |
| --- | --- | --- | --- |
|  | 1 | 2 | 3 |
|  | Lowest Capacity | Lowest Capacity | Lowest Capacity |
| 0 | 9782 | 9534 | 9263 |
| 1 | 7962 | 7899 | 7752 |

Figures 12 and 13 demonstrate how our proposed MILP model (19)-(22) for scenario 2 was able to determine the lowest capacity required for each site under 3 GDU possibilities while successfully schedule planting and harvesting weeks that resulted in consistent harvest quantities.

**Figure 12:**
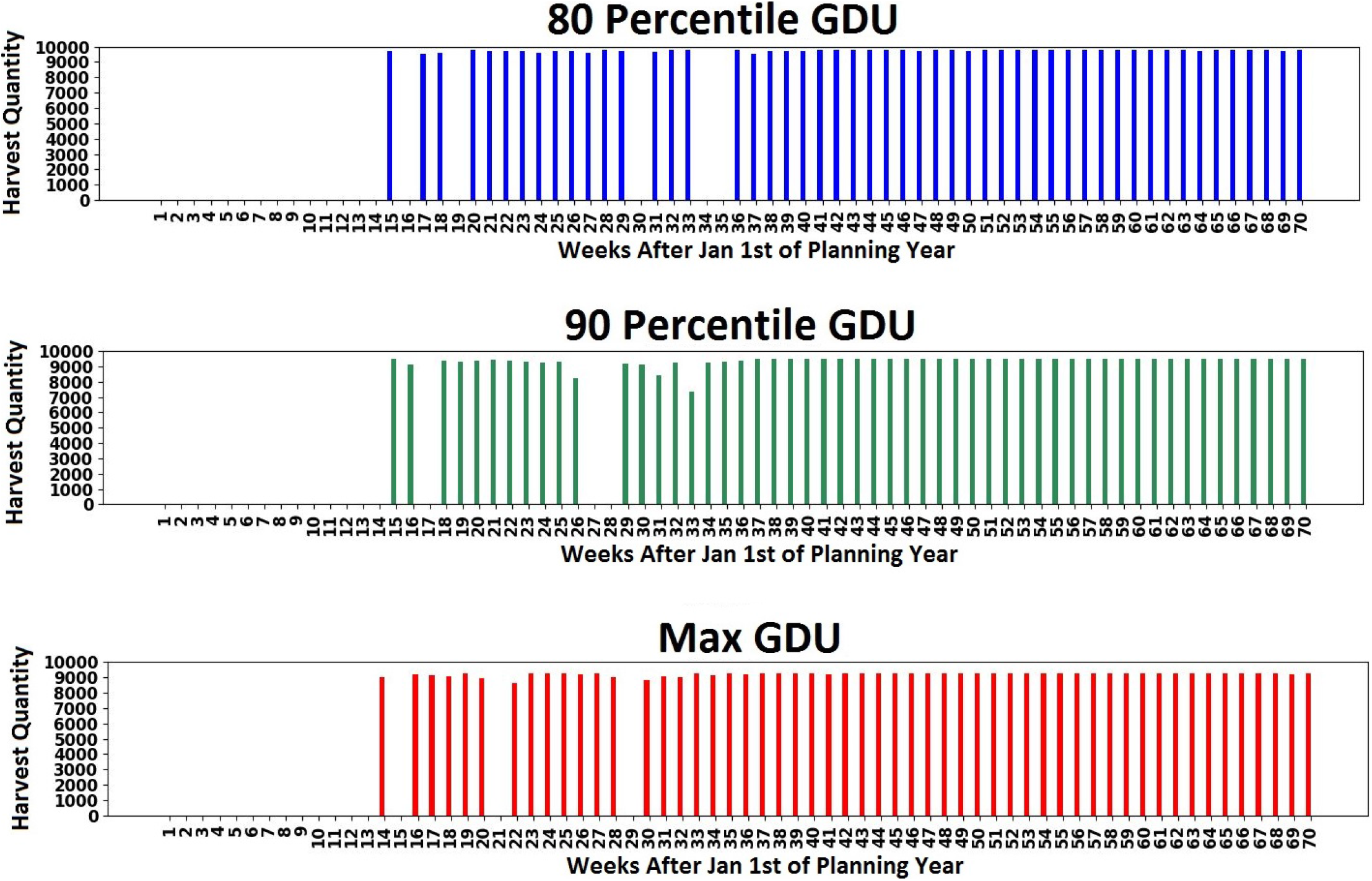
Weekly harvest quantity of 1375 seed populations planted in site 0 with suggested optimal capacity of 9782, 9534, and 9263 ears for 80 percentile, 90 percentile, and maximum GDU possibilities respectively for scenario 2.

**Figure 13:**
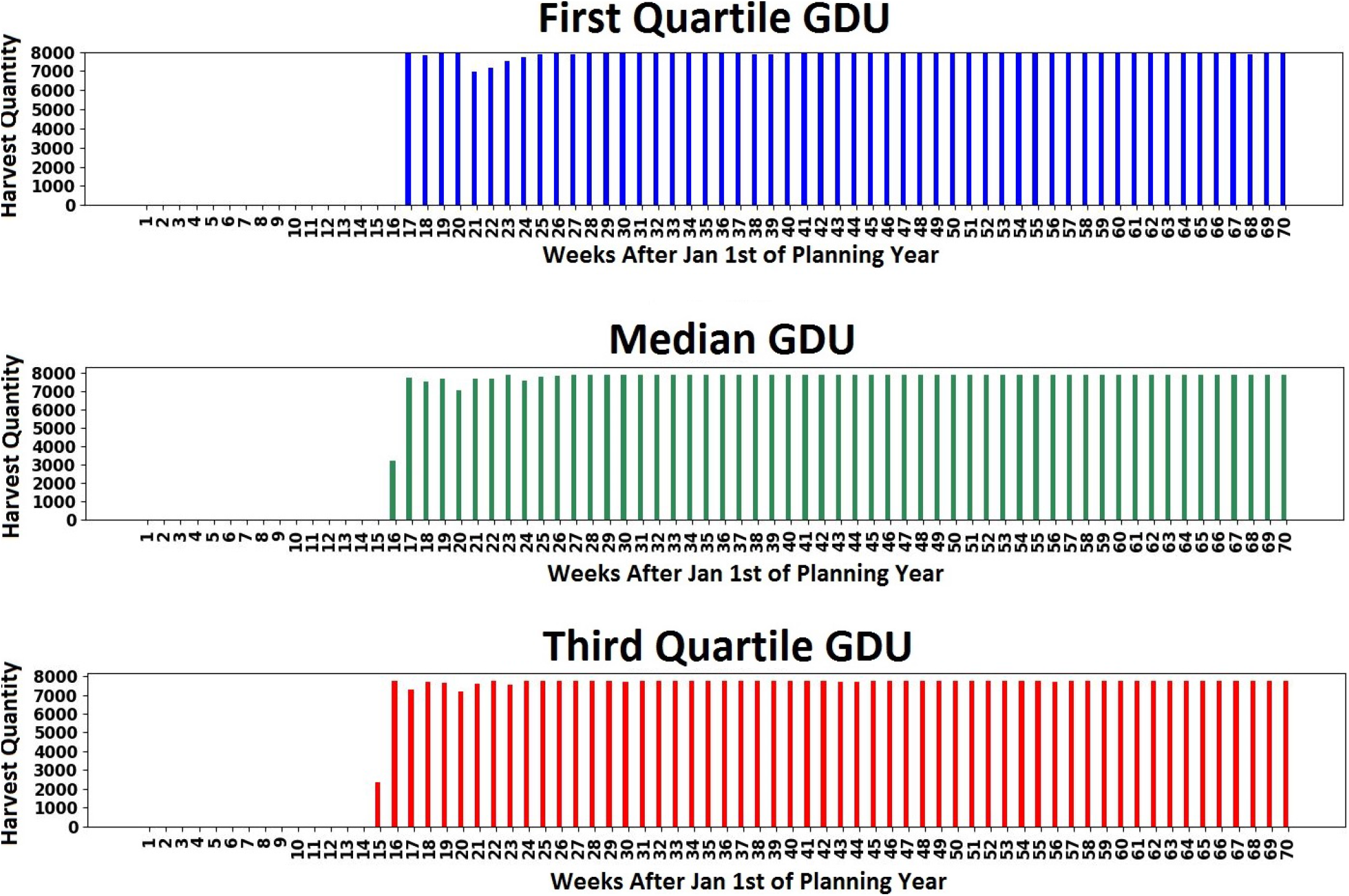
Weekly harvest quantity of 1194 seed populations planted in site 0 with suggested optimal capacity of 7962, 7899, and 7752 ears for first quartile, median, and third quartile GDU possibilities respectively for scenario 2.

Optimal planting and harvesting weeks for 25 seed populations from 1375 seed populations planted in site 0 and 1194 seed populations planted in site 1 are shown for each site under 3 GDU possibilities in figures 14 and 15.

**Figure 14:**
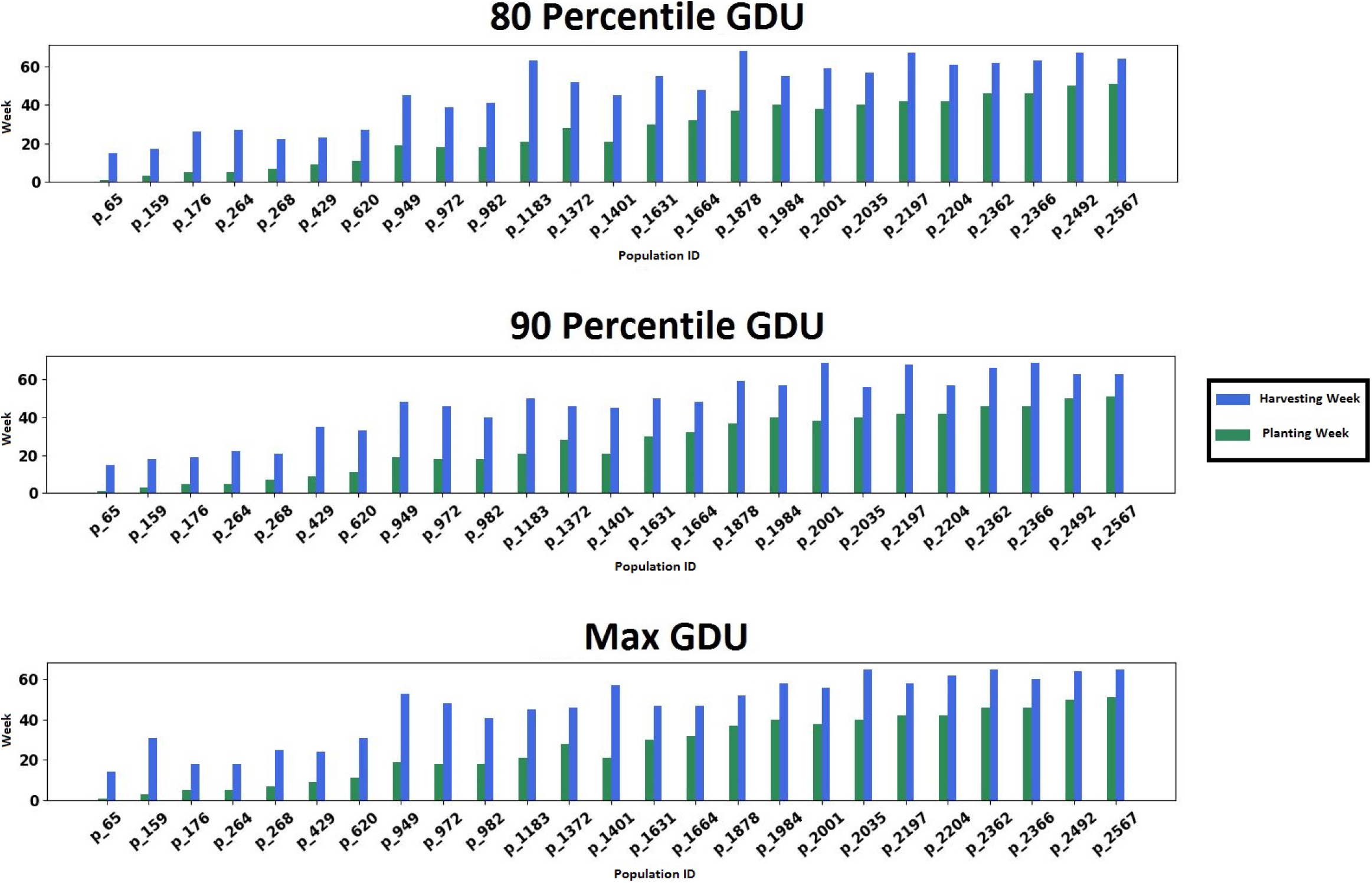
Optimal planting and harvesting week for a subset from the whole 1375 seed populations planted in site 0 under 3 GDU possibilities for scenario 2

**Figure 15:**
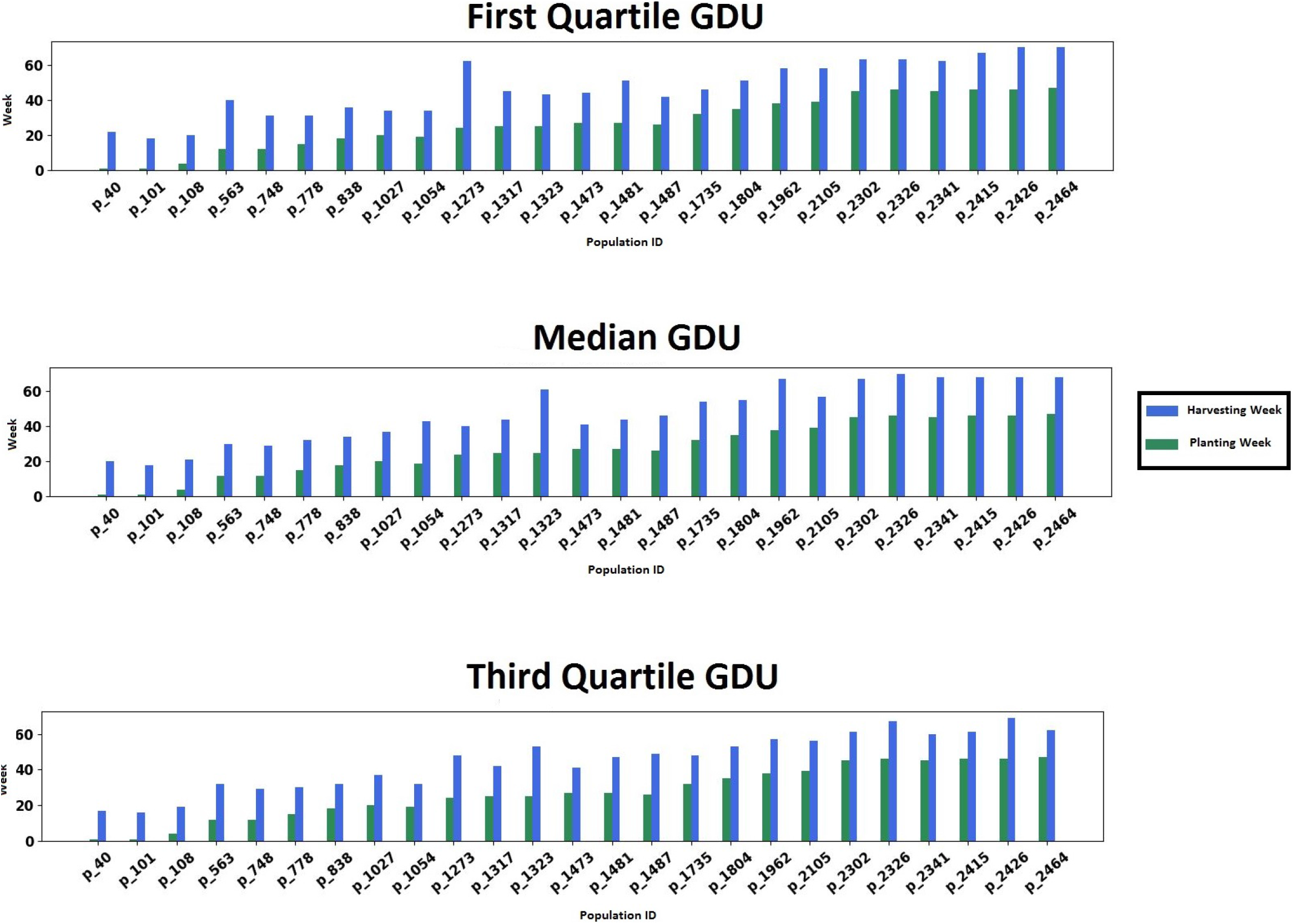
Optimal planting and harvesting week for a subset from the whole 1194 seed populations planted in site 1 under 3 GDU possibilities for scenario 2

## 6 Conclusion

In this paper, we proposed two MILP models for two storage capacity scenarios to help growers and farmers schedule planting and harvesting dates of different corn populations to have consistent harvest quantities that are below the capacity. Running our MILP models for different amounts of GDUs shows that the low average temperature or low daily GDUs makes the optimization models infeasible and prevent us from harvesting the whole populations in 70 weeks as the corn populations can not accumulate their required GDU to reach full maturity. Moreover, the results of our proposed MILP models indicate that different weather conditions or GDU quantities affect the number of harvesting weeks and harvest quantities. These explain why we determined the lowest GDU possibility and based on that we considered 3 different GDU possibilities for each site. Additionally, the results from MILP model for scenario 2 reveal that the amount of GDU units or weather condition also affect the lowest capacity required and the lower GDU units (lower average temperature) resulted in higher capacity required. It is because the corn populations accumulate their required GDUs slower, and as a result, higher number of corn populations should be harvested in later weeks. Therefore, higher storage capacity is required due to the limited number of harvesting weeks.

## Acknowledgements

We thank Syngenta and the Analytics Society of INFORMS for organizing the Syngenta Crop Challenge and providing the valuable datasets.

